# The genetic prehistory of the Greater Caucasus

**DOI:** 10.1101/322347

**Authors:** Chuan-Chao Wang, Sabine Reinhold, Alexey Kalmykov, Antje Wissgott, Guido Brandt, Choongwon Jeong, Olivia Cheronet, Matthew Ferry, Eadaoin Harney, Denise Keating, Swapan Mallick, Nadin Rohland, Kristin Stewardson, Anatoly R. Kantorovich, Vladimir E. Maslov, Vladimira G. Petrenko, Vladimir R. Erlikh, Biaslan Ch. Atabiev, Rabadan G. Magomedov, Philipp L. Kohl, Kurt W. Alt, Sandra L. Pichler, Claudia Gerling, Harald Meller, Benik Vardanyan, Larisa Yeganyan, Alexey D. Rezepkin, Dirk Mariaschk, Natalia Berezina, Julia Gresky, Katharina Fuchs, Corina Knipper, Stephan Schiffels, Elena Balanovska, Oleg Balanovsky, Iain Mathieson, Thomas Higham, Yakov B. Berezin, Alexandra Buzhilova, Viktor Trifonov, Ron Pinhasi, Andrej B. Belinskiy, David Reich, Svend Hansen, Johannes Krause, Wolfgang Haak

## Abstract

Archaeogenetic studies have described the formation of Eurasian ‘steppe ancestry’ as a mixture of Eastern and Caucasus hunter-gatherers. However, it remains unclear when and where this ancestry arose and whether it was related to a horizon of cultural innovations in the 4^th^ millennium BCE that subsequently facilitated the advance of pastoral societies likely linked to the dispersal of Indo-European languages. To address this, we generated genome-wide SNP data from 45 prehistoric individuals along a 3000-year temporal transect in the North Caucasus. We observe a genetic separation between the groups of the Caucasus and those of the adjacent steppe. The Caucasus groups are genetically similar to contemporaneous populations south of it, suggesting that – unlike today – the Caucasus acted as a bridge rather than an insurmountable barrier to human movement. The steppe groups from Yamnaya and subsequent pastoralist cultures show evidence for previously undetected farmer-related ancestry from different contact zones, while Steppe Maykop individuals harbour additional Upper Palaeolithic Siberian and Native American related ancestry.

---

The 1100-kilometre long Caucasus mountain ranges extend between the Black Sea and the Caspian Sea and are bound by the rivers Kuban and Terek in the north and by the Kura and Araxes rivers in the south. With Mount Elbrus in Russian Kabardino-Balkaria rising to a height of 5642 metres and Mount Shkhara in Georgia to 5201 metres, the Caucasus mountain ranges form a natural barrier between the Eurasian steppes and the Near East (Fig. 1).

**Fig. 1.**
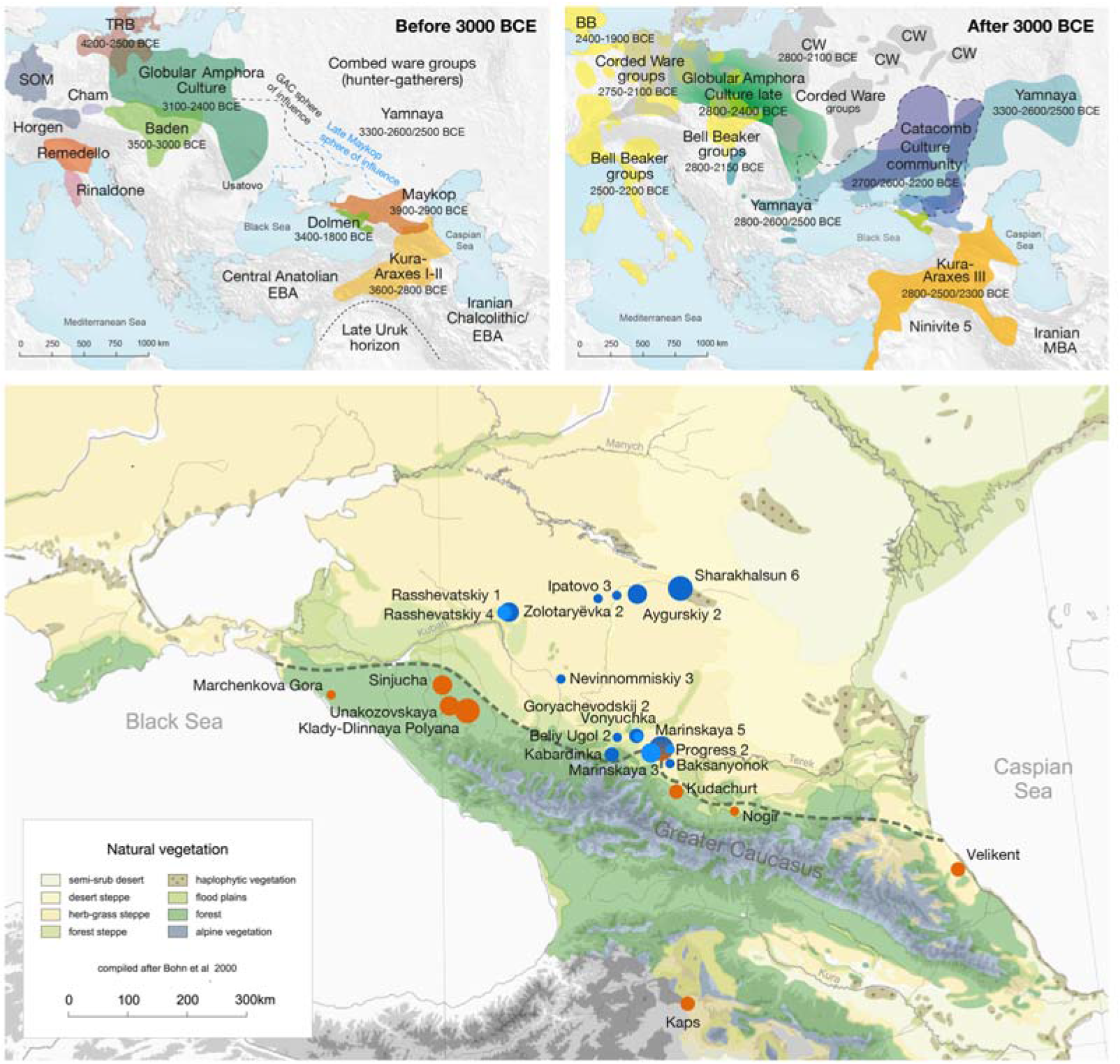
Map of sites and archaeological cultures mentioned in this study. Temporal and geographic distribution of archaeological cultures are shown for two windows in time that are critical for our data. The zoomed map shows the location of sites in the Caucasus. The size of the circle reflects number of individuals that produced genome-wide data. The dashed line illustrates a hypothetical geographic border between genetically distinct *Steppe* and *Caucasus* clusters. *(BB=Bell Beaker; CW=Corded Ware; TRB=Trichterbecher/Funnel Beaker; SOM=Seine-Oise-Marne complex)*

The rich archaeological record suggests extensive periods of human occupation since the Upper Palaeolithic^1, 2, 3^. The density of languages and cultures in the region is mirrored by faunal and floral diversity, and the Caucasus has often been described as a contact zone and natural refuge with copious ecological niches. However, it also serves as a bio-geographic border between the steppe and regions to the south such as Anatolia and Mesopotamia rather than a corridor for human^4, 5^ and animal movement^6, 7, 8^. The extent to which the Caucasus has played an important role for human population movements between south and north over the course of human history is thus a critical question, and one that until now has been unanswered by archaeogenetic studies.

A Neolithic lifestyle based on food production began in the Caucasus after 6000 calBCE^9^. In the following millennia the Caucasus region began to play an increasingly important role in the economies of the growing urban centres in northern Mesopotamia^10^ as a region rich in natural resources such as ores, pastures and timber^11^. In the 4^th^ millennium BCE the archaeological record attests to the presence of the Maykop and Kura-Araxes cultural complexes, with the latter being found on both flanks of the Caucasus mountain range, thus clearly demonstrating the connection between north and south^11^. The Maykop culture was an important player in the innovative horizon of the 4^th^ millennium BCE in Western Eurasia. It is well known for its rich burial mounds, especially at the eponymous Maykop site in today‘s Adygea, which reflect the rise of a new system of social organization^12^. The 4^th^ millennium BCE witnesses a concomitant rise in commodities and technologies such as the wheel and wagon including associated technology, copper alloys, new weaponry, and new breeds of domestic sheep^13, 14^.

The adjacent Pontic-Caspian and Eurasian steppe also played an important role in this linked economic system, being the most likely region for the domestication of the horse that revolutionised transport^13^. In addition, many steppe kurgans (large burial mounds that are first observed in the context of the Maykop culture) have yielded the remains of wheels and ox-drawn carts, highlighting a mobile economy focused on cattle and sheep/goat herding^15^. The adoption of the horse almost certainly contributed to the intensification of pastoralist practices in the Eurasian steppes, allowing more efficient keeping of larger herds^16, 17, 18^ and facilitating the massive range expansions of pastoralists associated with the Yamnaya cultural community and related groups from the East European steppe^19, 20^. This transformation changed the European gene pool during the early 3^rd^millennium BCE and descendants of the Yamnaya eventually also transformed the ancestry of South Asia as well^21^. However, flow of goods and ideas between the eastern European steppe zone, the Caucasus, the Carpathians, and Central Europe has been documented by archaeological and ancient DNA research as early as the 5^th^ millennium BCE, long before the massive migration took place^22, 23, 24^. Taken together, the Caucasus region played a crucial role in the prehistory of Western Eurasia and this study aims to shed new light on events in the key period between the 4^th^ and 3^rd^ millennium BCE.

Recent ancient DNA studies have enabled the resolution of several long-standing questions regarding cultural and population transformations in prehistory. One of these is the Mesolithic-Neolithic transition in Europe, which saw a change from a hunter-gatherer lifestyle to a sedentary, food-producing subsistence strategy. Genome-wide data from pre-farming and farming communities have identified distinct ancestral populations that largely reflect subsistence patterns in addition to geography^25^. One important feature is a cline of European hunter-gatherer (HG) ancestry that runs roughly from West to East (hence WHG and EHG; blue component in Fig. 2A, 2C), which differs greatly from the ancestry of Early European farmers that in turn is closely related to that of northwest Anatolian farmers^26, 27^ and more remotely also to pre-farming individuals from the Levant^23^. The Near East and Anatolia have long been seen as the regions from which European farming and animal husbandry emerged. Surprisingly, these regions harboured three divergent populations, with Anatolian and Levantine ancestry in the western part and a group with a distinct ancestry in the eastern part first described in Upper Pleistocene individuals from Georgia (Caucasus hunter-gatherers; CHG)^2^ and then in Mesolithic and Neolithic individuals from Iran^23, 29^. The following two millennia, spanning from the Neolithic to Chalcolithic and Early Bronze Age periods in each region, witnessed migration and admixture between these ancestral groups, leading to a pattern of genetic homogenization and reduced genetic distances between these Neolithic source populations^23^. In parallel, Eneolithic individuals from the Samara region (5200-4000 BCE) also exhibit population mixture, specifically EHG-and CHG/Iranian ancestry, a combination that forms the so-called ‘steppe-ancestry’^28^. This ancestry eventually spread further west^19, 20^, where it contributed substantially to the ancestry of present-day Europeans, and east to the Altai region as well as to South Asia^23^.

**Fig. 2.**
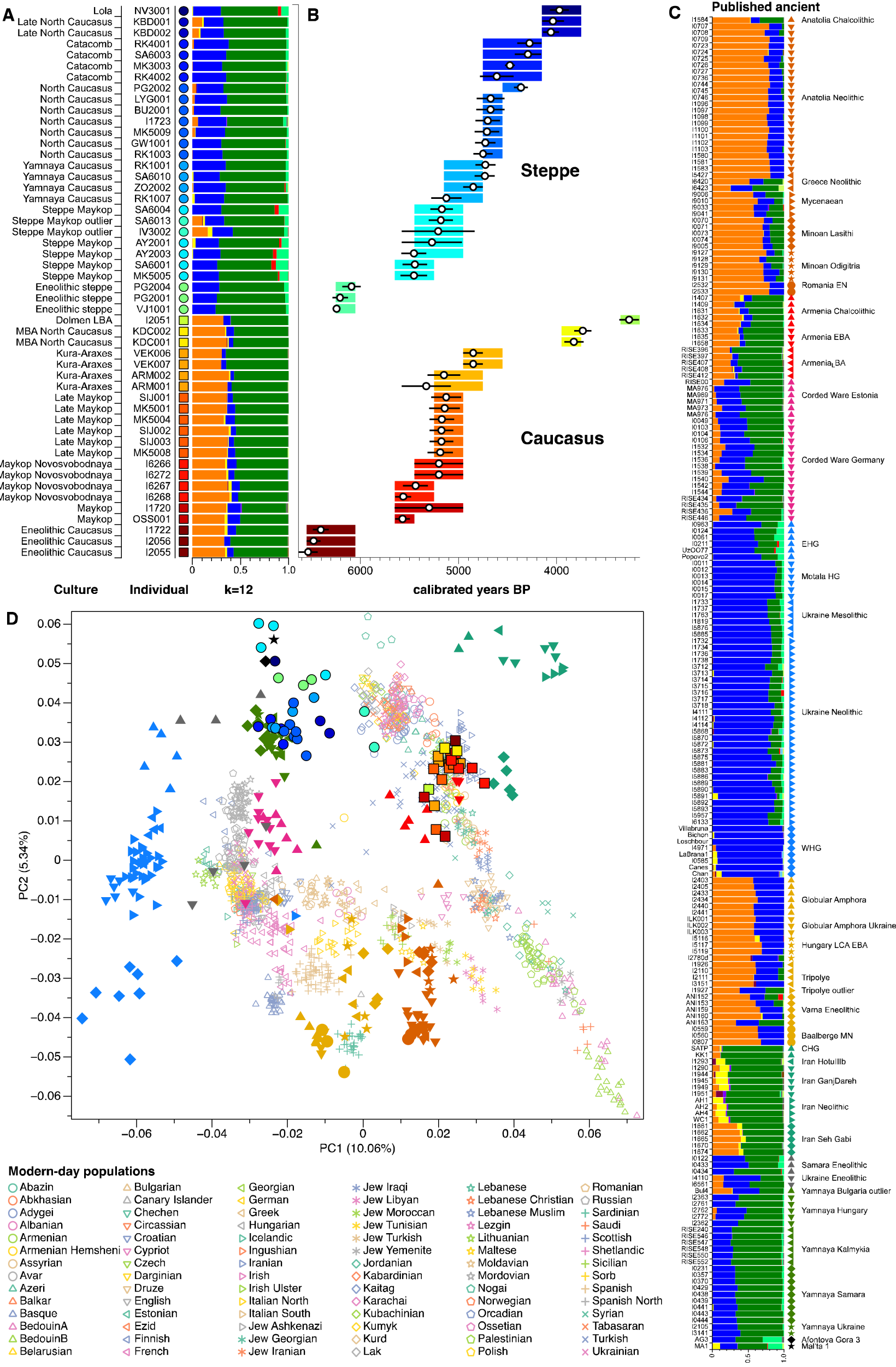
ADMIXTURE and PCA results, and chronological order of ancient Caucasus individuals. (a) *ADMIXTURE* results (k=12) of the newly genotyped individuals (filled symbols with black outlines) sorted by genetic clusters (*Steppe* and *Caucasus*) and in chronological order(coloured bars indicate the relative archaeological dates, (b) white circles the mean calibrated radiocarbon date and the errors bars the 2-sigma range. (c) *ADMIXTURE* results of relevant prehistoric individuals mentioned in the text (filled symbols) and (d) shows these projected onto a PCA of 84 modern-day West Eurasian populations (open symbols).

To understand and characterize the genetic variation of Caucasian populations, present-day groups from various geographic, cultural/ethnic and linguistic backgrounds have been analyzed previously at the autosomal, Y-chromosomal and mitochondrial level^4, 5, 30^. Yunusbayev and colleagues described the Greater Caucasus region as an asymmetric semipermeable barrier based on a higher genetic affinity of southern Caucasus groups to Anatolian and Near Eastern populations and a genetic discontinuity between these and populations of the North Caucasus and of adjacent Eurasian steppes. While autosomal and mitochondrial DNA data appear relatively homogeneous across diverse ethnic and linguistic groups and the entire mountainous region, the Y-chromosome diversity reveals a deeper genetic structure attesting to several male founder effects, with striking correspondence to geography, language groups and historical events^4, 5^.

In our study we aimed to investigate when and how the genetic patterns observed today were formed and test whether they have been present since prehistoric times by generating time-stamped human genome-wide data. We were also interested in characterizing the role of the Caucasus as a conduit for gene-flow in the past and in shaping the cultural and genetic makeup of the wider region (Supplementary Information 1). This has important implications for understanding the means by which Europe, the Eurasian steppe zone, and the earliest urban centres in the Near East were connected^31^. We aimed to genetically characterise individuals from cultural complexes such as the Maykop and Kura-Araxes and assessing the amount ofgene flow in the Caucasus during times when the exploitation of resources of the steppe environment intensified, since this was potentially triggered by the cultural and technological innovations of the Late Chalcolithic and Early Bronze Age 6000-5000 years ago^11^. Lastly, since the spread of steppe ancestry into central Europe and the eastern steppes during the early 3^rd^ millennium BCE (5000-4500 BP) was a striking migratory event in human prehistory^19, 20^, we also wanted to retrace the formation of the steppe ancestry profile and whether this might have been influenced by neighbouring farming groups to the west or from regions of early urbanization further south.

## Results

### Genetic clustering and uniparentally-inherited markers

We report genome-wide data at a targeted set of 1.2 million single nucleotide polymorphisms (SNPs)^19, 32^ for 59 Eneolithic/Chalcolithic and Bronze Age individuals from the Caucasus region. After filtering out 14 individuals that were first-degree relatives or showed evidence of contamination or reference bias (Supplementary Information 3 and Data 1) we retained 45 individuals for downstream analyses using a cut-off of 30,000 SNPs. We merged our newly generated samples with previously published ancient and modern data^19, 20, 23, 24, 26, 27 29, 33, 34, 35, 36, 37 38, 39,40, 41, 42, 43^ (Supplementary Data 2). We first performed principal component analysis (PCA)^44^ and ADMIXTURE^45^ analysis to assess the genetic affinities of the ancient individuals qualitatively (Fig. 2) and followed up quantitatively with formal *f*- and *D*- statistics, *qpWave, qpAdm,* and *qpGraph^44^.* Based on PCA and ADMIXTURE plots we observe two distinct genetic clusters: one cluster falls with previously published ancient individuals from the West Eurasian steppe (hence termed ‘*Steppe*’), and the second clusters with present-day southern Caucasian populations and ancient Bronze Age individuals from today’s Armenia (henceforth called ‘*Caucasus*’), while a few individuals take on intermediate positions between the two. The stark distinction seen in our temporal transect is also visible in the Y-chromosome haplogroup distribution, with R1/R1b1 and Q1a2 types in the *Steppe* and L, J, and G2 types in the *Caucasus* cluster (Fig. 3A, Supplementary Data 1). In contrast, the mitochondrial haplogroup distribution is more diverse and almost identical in both groups (Fig. 3B, Supplementary Data 1).

**Fig. 3.**
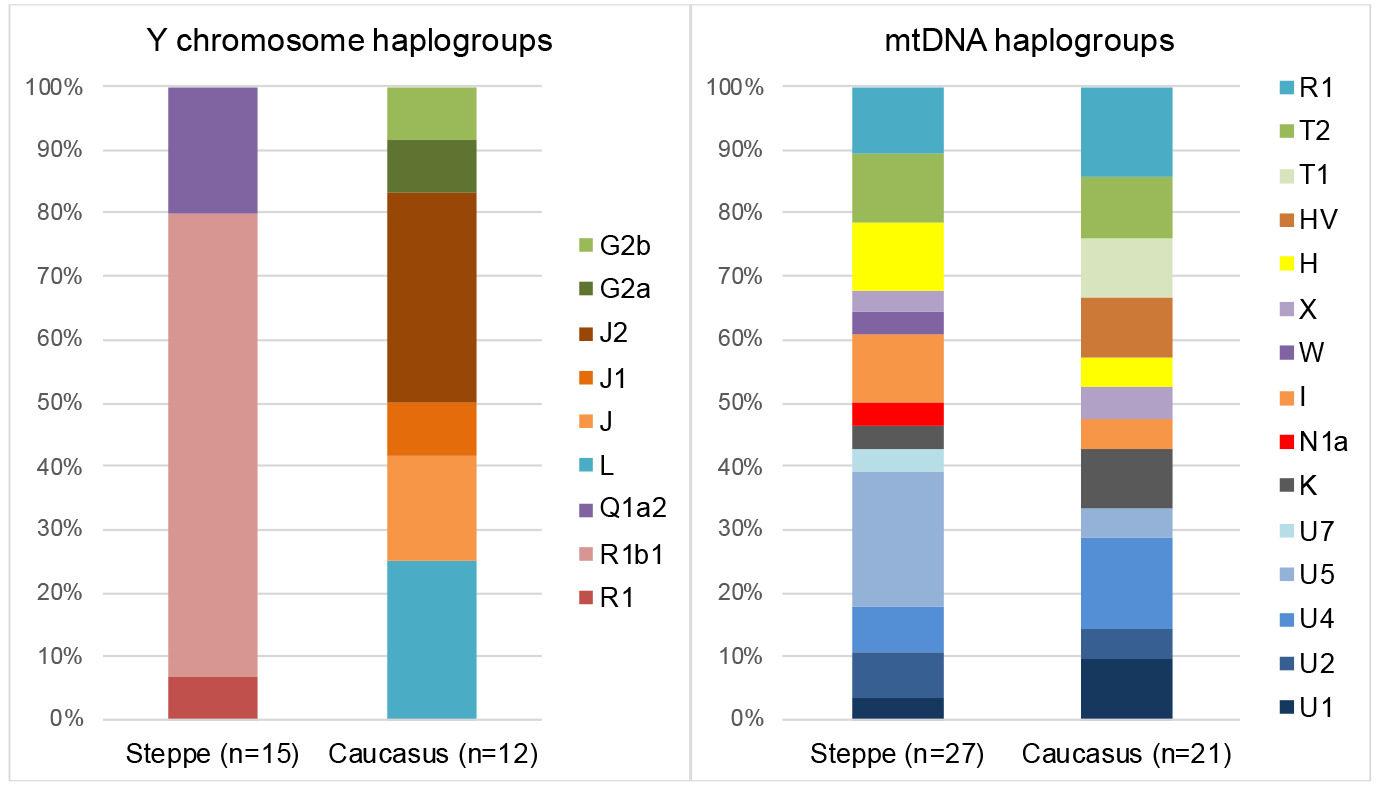
Comparison of Y-chromosome (A) and mitochondrial (B) haplogroup distribution in the *Steppe* and *Caucasus* cluster.

The two distinct clusters are already visible in the oldest individuals of our temporal transect, dated to the Eneolithic period (~6300-6100 yBP/4300-4100 calBCE). Three individuals from the sites of Progress 2 and Vonjuchka 1 in the North Caucasus piedmont steppe (‘Eneolithic steppe’), which harbor Eastern and Caucasian hunter-gatherer related ancestry (EHG and CHG, respectively), are genetically very similar to Eneolithic individuals from Khalynsk II and the Samara region^19, 27^ This extends the cline of dilution of EHG ancestry via CHG/Iranian-like ancestry to sites immediately north of the Caucasus foothills (Fig. 2D).

In contrast, the oldest individuals from the northern mountain flank itself, which are three first degree-related individuals from the Unakozovskaya cave associated with the Darkveti-Meshoko Eneolithic culture (analysis label ‘Eneolithic Caucasus’) show mixed ancestry mostly derived from sources related to the Anatolian Neolithic (orange) and CHG/Iran Neolithic (green) in the ADMIXTURE plot (Fig. 2C). While similar ancestry profiles have been reported for Anatolian and Armenian Chalcolithic and Bronze Age individuals^20, 23^, this result suggests the presence of the mixed Anatolian/Iranian/CHG related ancestry north of the Great Caucasus Range as early as ~6500 years ago.

### Ancient North Eurasian ancestry in ‘Steppe Maykop’ individuals

Four individuals from mounds in the grass steppe zone, which are archaeologically associated with the ‘Steppe Maykop’ cultural complex (Supplementary Information 1), lack the Anatolian farmer-related component when compared to contemporaneous Maykop individuals from the foothills. Instead they carry a third and fourth ancestry component that is linked deeply to Upper Paleolithic Siberians (maximized in the individual Afontova Gora 3 (AG3)^36, 37^ and Native Americans, respectively, and in modern-day North Asians such as North Siberian Nganasan (Supplementary Fig. 1). To illustrate this affinity with ‘ancient North Eurasians’ (ANE)^26^, we also ran PCA with 147 Eurasian (Supplementary Fig. 2A) and 29 Native American populations (Supplementary Fig. 2B). The latter represent a cline from ANE-rich steppe populations such as EHG, Eneolithic individuals, AG3 and Mal’ta 1 (MA1) to modern-day Native Americans at the opposite end. To formally test the excess of alleles shared with ANE/Native Americans we performed *f_4_*-statistics of the form *f_4_*(Mbuti, X; Steppe Maykop, Eneolithic steppe), which resulted in significantly positive Z scores |Z >3| for AG3, MA1, EHG, Clovis and Kennewick for the ancient populations and many present-day Native American populations (Supplementary Table 1). Based on these observations we used *qpWave* and *qpAdm* methods to model the number of ancestral sources contributing to the Steppe Maykop individuals and their relative ancestry coefficients. Simple two-way models of Steppe Maykop as an admixture of Eneolithic steppe, AG3 or Kennewick do not fit (Supplementary Table 2). However, we could successfully model Steppe Maykop ancestry as being derived from populations related to all three sources (p-value 0.371 for rank 2): Eneolithic steppe(63.5±2.9 %), AG3 (29.6±3.4%) and Kennewick (6.9±1.0%) (Fig. 4; Supplementary Table 3). We note that the Kennewick related signal is most likely driven by the East Eurasian part of Native American ancestry as the *f_4_*statistics (Steppe_Maykop, Fitted Steppe_Maykop; Outgroup1, Outgroup2) show that the Steppe Maykop individuals share more alleles not only with Karitiana but also with Han Chinese when compared with the fitted ones using Eneolithic steppe and AG3 as two sources and Mbuti, Karitiana and Han as outgroups (Supplementary Table 2).

**Fig. 4.**
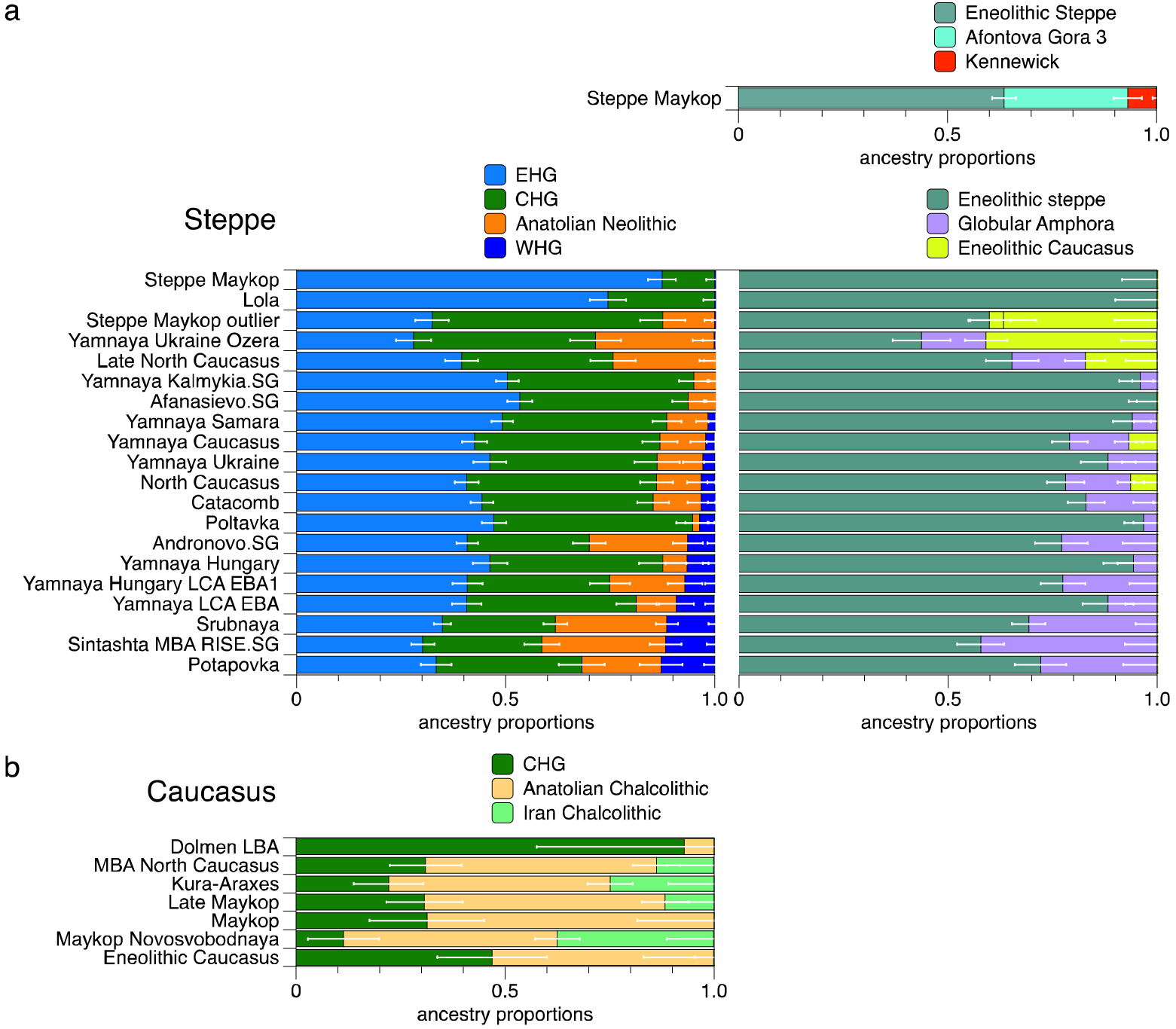
Modelling results for the Steppe and Caucasus cluster. Admixture proportions based on (temporally and geographically) distal and proximal models, showing additional Anatolian farmer-related ancestry in Steppe groups as well as additional gene flow from the south in some of the Steppe groups as well as the Caucasus groups (see also Supplementary Tables 10, 14 and 20).

### Characterising the *Caucasus* ancestry profile

The Maykop period, represented by twelve individuals from eight Maykop sites (Maykop, n=2; a cultural variant ‘Novosvobodnaya’ from the site Klady, n=4; and Late Maykop, n=6) in the northern foothills appear homogeneous. These individuals closely resemble the preceding Caucasus Eneolithic individuals and present a continuation of the local genetic profile. This ancestry persists in the following centuries at least until ~3100 yBP (1100 calBCE) in the mountains, as revealed by individuals from Kura-Araxes from both the northeast (Velikent, Dagestan) and the South Caucasus (Kaps, Armenia), as well as Middle and Late Bronze Age individuals (e.g. Kudachurt, Marchenkova Gora) from the north. Overall, this *Caucasus* ancestry profile falls among the ‘Armenian and Iranian Chalcolithic’ individuals and is indistinguishable from other Kura-Araxes individuals (‘Armenian Early Bronze Age’) on the PCA plot (Fig. 2), suggesting a dual origin involving Anatolian/Levantine and Iran Neolithic/CHG ancestry, with only minimal EHG/WHG contribution possibly as part of the Anatolian farmer-related ancestry^23^.

Admixture *f_3_*−statistics of the form *f_3_*(X, Y; target) with the *Caucasus* cluster as target resulted in significantly negative Z scores |Z < −3| when CHG (or AG3 in Late Maykop) were used as one and Anatolian farmers as the second potential source (Supplementary Table 4). We also used *qpWave* to determine the number of streams of ancestry and found that a minimum of two is sufficient (except for Eneolithic Caucasus or Dolmen LBA, for which one source is sufficient (Supplementary Table 5).

We then tested whether each temporal/cultural group of the *Caucasus* cluster could be modelled as a simple two-way admixture by exploring all possible pairs of sources in *qpWave*. We found support for CHG as one source and Anatolian farmer-related ancestry or a derived form such as is found in southeastern Europe as the other (Supplementary Table 6). We focused on model of mixture of proximal sources (Fig. 4B) such as CHG and Anatolian Chalcolithic for all six groups of the Caucasus cluster (Eneolithic Caucasus, Maykop and Late Makyop, Maykop-Novosvobodnaya, Kura-Araxes, and Dolmen LBA), with admixture proportions on a genetic cline of 40-72% Anatolian Chalcolithic related and 28-60% CHG related (Supplementary Table 7). When we explored Romania_EN and Greece_Neolithic individuals as alternative southeast European sources (30-46% and 36-49%), the CHG proportions increased to 54-70% and 51-64%, respectively. We hypothesize that alternative models, replacing the Anatolian Chalcolithic individual with yet unsampled populations from eastern Anatolia, South Caucasus or northern Mesopotamia, would probably also provide a fit to the data from some of the tested Caucasus groups. The models replacing CHG with Iran Neolithic-related individuals could explain the data in a two-way admixture with the combination of Armenia Chalcolithic or Anatolia Chalcolithic as the other source. However, models replacing CHG with EHG individuals received no support (Supplementary Table 8), indicating no strong influence for admixture from the adjacent steppe to the north. In an attempt to account for potentially un-modelled ancestry in the Caucasus groups, we added EHG, WHG and Iran Chalcolithic as additional sources in the previous two-way modelling. The resulting ancestry coefficients do not deviate substantially from 0 (high standard errors) when adding EHG or WHG, suggesting very limited direct ancestry from both hunter-gatherer groups (Supplementary Table 9). Alternatively, when we added Iran Chalcolithic individuals as a third source to the model, we observed that Kura-Araxes and Maykop-Novosvobodnaya individuals had likely received additional Iran Chalcolithic-related ancestry (24.9% and 37.4%, respectively; Fig. 4; Supplementary Table 10).

### Characterising the *Steppe* ancestry profile in the North Caucasus

Individuals from the North Caucasian steppe associated with the Yamnaya cultural formation (5300-4400 BP, 3300-2400 calBCE) appear genetically almost identical to previously reported Yamnaya individuals from Kalmykia^20^ immediately to the north, the middle Volga region^19, 27^, Ukraine and Hungary, and to other Bronze Age individuals from the Eurasian steppes who share the characteristic ‘steppe ancestry’ profile as a mixture of EHG and CHG/Iranian ancestry^23, 28^. These individuals form a tight cluster in PCA space (Figure 2) and can be shown formally to be a mixture by significantly negative admixture *f_3_*statistics of the form *f_3_*(EHG, CHG; target) (Supplementary Fig. 3). This also involves individuals assigned to the North Caucasus culture (4800-4500 BP, 2800-2500 calBCE) in the piedmont steppe of the central North Caucasus, who share the steppe ancestry profile. Individuals from the Catacomb culture in the Kuban, Caspian and piedmont steppes (4600-4200 BP, 2600-2200 calBCE), which succeeded the Yamnaya horizon, also show a continuation of the ‘steppe ancestry’ profile.

The individuals of the Middle Bronze Age (MBA) post-Catacomb horizon (4200-3700 BP,2200-1700 calBCE) such as Late North Caucasus and Lola culture represent both ancestry profiles common in the North Caucasus region: individuals from the mountain site Kabardinka show a typical steppe ancestry profile, whereas individuals from the Late North Caucasus site Kudachurt 90 km to the west retain the ‘southern’ Caucasus profile. The latter is also observed in our most recent individual from the western Late Bronze Age Dolmen culture (3400-3200 BP, 1400-1200 calBCE). In contrast, one individual assigned to the Lola culture resembles the ancestry profile of the Steppe Maykop individuals.

### Admixture into the steppe zone from the south

Evidence for interaction between the *Caucasus* and the *Steppe* clusters is visible in our genetic data from individuals associated with the later Steppe Maykop phase around 5300-5100 years ago. These ‘outlier’ individuals were buried in the same mounds as those with steppe and in particular Steppe Maykop ancestry profiles but share a higher proportion of Anatolian farmer-related ancestry visible in the ADMIXTURE plot and are also shifted towards the *Caucasus* cluster in PC space (Fig. 2D). This observation is confirmed by formal *D*-statistics (Steppe Maykop outlier, Steppe Maykop; X; Mbuti), which are significantly positive when X is a Neolithic or Bronze Age group from the Near East or Anatolia (Supplementary Fig.4). By modelling Steppe Maykop outliers successfully as a two-way mixture of Steppe Maykop and representatives of the *Caucasus* cluster (Supplementary Table 3), we can show that these individuals received additional ‘Anatolian and Iranian Neolithic ancestry’, most likely from contemporaneous sources in the south. We estimated admixture time for the observed farmer-related ancestry individuals using the linkage disequilibrium (LD)-based admixture inference implemented in ALDER^46^, using Steppe Maykop outliers as the test population and Steppe Maykop as well as Kura-Araxes as references. The average admixture time for Steppe Maykop outliers is about 20 generations or 560 years ago, assuming a generation time of 28 years^47^ (Supplementary Information 6).

### Contribution of Anatolian farmer-related ancestry to Bronze Age steppe groups

In principal component space Eneolithic individuals (Samara Eneolithic) form a cline running from EHG to CHG (Fig. 2D), which is continued by the newly reported Eneolithic steppe individuals. However, the trajectory of this cline changes in the subsequent centuries. Here we observe a cline from Eneolithic_steppe towards the *Caucasus* cluster. We can qualitatively explain this ‘tilting cline’ by developments south of the Caucasus, where Iranian and Anatolian/Levantine Neolithic ancestries continue to mix, resulting in a blend that is also observed in the *Caucasus* cluster, from where it could have spread onto the steppe. The first appearance of ‘Near Eastern farmer related ancestry’ in the steppe zone is evident in Steppe Maykop outliers. However, PCA results also suggest that Yamnaya and later groups of the West Eurasian steppe carry some farmer related ancestry as they are slightly shifted towards ‘European Neolithic groups’ in PC2 (Fig. 2D) compared to Eneolithic steppe. This is not the case for the preceding Eneolithic steppe individuals. The tilting cline is also confirmed by admixture *f_3_*-statistics, which provide statistically negative values for AG3 as one source and any Anatolian Neolithic related group as a second source (Supplementary Table 11). Detailed exploration via *D*-statistics in the form of *D*(EHG, steppe group; X, Mbuti) and *D*(Samara_Eneolithic, steppe group; X, Mbuti) show significantly negative *D* values for most of the steppe groups when X is a member of the *Caucasus* cluster or one of the Levant/Anatolia farmer-related groups (Supplementary Figs. 5 and 6). In addition, we used *f*- and *D*-statistics to explore the shared ancestry with Anatolian Neolithic as well as the reciprocal relationship between Anatolian- and Iranian farmer-related ancestry for all groups of our two main clusters and relevant adjacent regions (Supplementary Fig. 4). Here, we observe an increase in farmer-related ancestry (both Anatolian and Iranian) in our *Steppe* cluster, ranging from Eneolithic steppe to later groups. In Middle/Late Bronze Age groups especially to the north and east we observe a further increase of Anatolian farmer-related ancestry consistent with previous studies of the Poltavka, Andronovo, Srubnaya and Sintashta groups^23, 27^ and reflecting a different process not especially related to events in the Caucasus.

The exact geographic and temporal origin of this Anatolian farmer-related ancestry in the North Caucasus and later in the steppe is difficult to discern from our data. Not only do the *Steppe* groups vary in their respective affinity to each of the two, but also the *Caucasus* groups, which represent potential sources from a geographic and cultural point of view, are mixtures of them both^23^. We therefore used *qpWave* and *qpAdm* to explore the number of ancestry sources for the Anatolian farmer-related component to evaluate whether geographically proximate groups plausibly contributed to the subtle shift of Eneolithic ancestry in the steppe towards those of the Neolithic groups. Specifically, we tested whether any of the Eurasian steppe ancestry groups can be successfully modelled as a two-way admixture between Eneolithic steppe and a population X derived from Anatolian-or Iranian farmer-related ancestry, respectively. Surprisingly, we found that a minimum of four streams of ancestry is needed to explain all eleven steppe ancestry groups tested, including previously published ones (Fig. 2; Supplementary Table 12). Importantly, our results show a subtle contribution of both Anatolian farmer-related ancestry and WHG-related ancestry (Fig.4; Supplementary Tables 13 and 14), which was likely contributed through Middle and Late Neolithic farming groups from adjacent regions in the West. A direct source of Anatolian farmer-related ancestry can be ruled out (Supplementary Table 15). At present, due to the limits of our resolution, we cannot identify a single best source population. However, geographically proximal and contemporaneous groups such as Globular Amphora and Eneolithic groups from the Black Sea area (Ukraine and Bulgaria), which represent all four distal sources (CHG, EHG, WHG, and Anatolian_Neolithic) are among the best supported candidates (Fig. 4; Supplementary Tables 13,14 and 15). Applying the same method to the subsequent North Caucasian *Steppe* groups such as Catacomb, North Caucasus, and Late North Caucasus confirms this pattern (Supplementary Table 17).

Using *qpAdm* with Globular Amphora as a proximate surrogate population (assuming that a related group was the source of the Anatolian farmer-related ancestry), we estimated the contribution of Anatolian farmer-related ancestry into Yamnaya and other steppe groups. We find that Yamnaya individuals from the Volga region (Yamnaya Samara) have 13.2±2.7% and Yamnaya individuals in Hungary 17.1±4.1% Anatolian farmer-related ancestry (Fig.4; Supplementary Table 18)– statistically indistinguishable proportions. Replacing Globular Amphora by Iberia Chalcolithic, for instance, does not alter the results profoundly (Supplementary Table 19). This suggests that the source population was a mixture of Anatolian farmer-related ancestry and a minimum of 20% WHG ancestry, a profile that is shared by many Middle/Late Neolithic and Chalcolithic individuals from Europe of the 3^rd^ millennium BCE analysed thus far.

To account for potentially un-modelled ancestry from the *Caucasus* groups, we added ‘Eneolithic Caucasus’ as an additional source to build a three-way model. We found that Yamnaya Caucasus, Yamnaya Ukraine Ozera, North Caucasus and Late North Caucasus had likely received additional ancestry (6% to 40%) from nearby *Caucasus* groups (Supplementary Table 20). This suggests a more complex and dynamic picture of steppe ancestry groups through time, including the formation of a local variant of steppe ancestry in the North Caucasian steppe from the local Eneolithic, a contribution of Steppe Maykop groups, and population continuity between the early Yamnaya period and the Middle Bronze Age (5300-3200 BP, 3300-2200 calBCE). This was interspersed by additional, albeit subtle gene-flow from the West and occasional equally subtle gene flow from neighbouring groups in the Caucasus and piedmont zones.

### Insights from micro-transects through time

The availability of multiple individuals from one site (here burial mounds or kurgans) allowed us to test genetic continuity on a micro-transect level. By focusing on two kurgans (Marinskaya 5 and Sharakhalsun 6), for which we could successfully generate genome-wide data from four and five individuals, respectively, we observe that the genetic ancestry varied through time, alternating between the *Steppe* and *Caucasus* ancestries (Supplementary Fig. 8). This shows that the apparent genetic border between the two distinct genetic clusters was shifting over time. We also detected various degrees of kinship between individuals buried in the same mound, which supports the view that particular mounds reflected genealogical lineages. Overall, we observe a balanced sex ratio within our sites across the individuals tested (Supplementary Information 4).

### A joint model of ancient populations of the Caucasus region

We used *qpGraph* to explore models that jointly explain the population splits and gene flow in the Greater Caucasus region by computing *f_2_*-, *f_3_*- and *f_4_*- statistics measuring allele sharing among pairs, triples, and quadruples of populations and evaluating fits based on the maximum |Z|-score comparing predicted and observed values of these statistics. Our fitted model recapitulates the genetic separation between the *Caucasus* and *Steppe* groups with the Eneolithic steppe individuals deriving more than 60% of ancestry from EHG and the remainder from a CHG-related basal lineage, whereas the Maykop group received about 86.4% from CHG, 9.6% Anatolian farming related ancestry, and4% from EHG. The Yamnaya individuals from the Caucasus derived the majority of their ancestry from Eneolithic steppe individuals but also received about 16% from Globular Amphora-related farmers (Fig. 5).

**Fig. 5.**
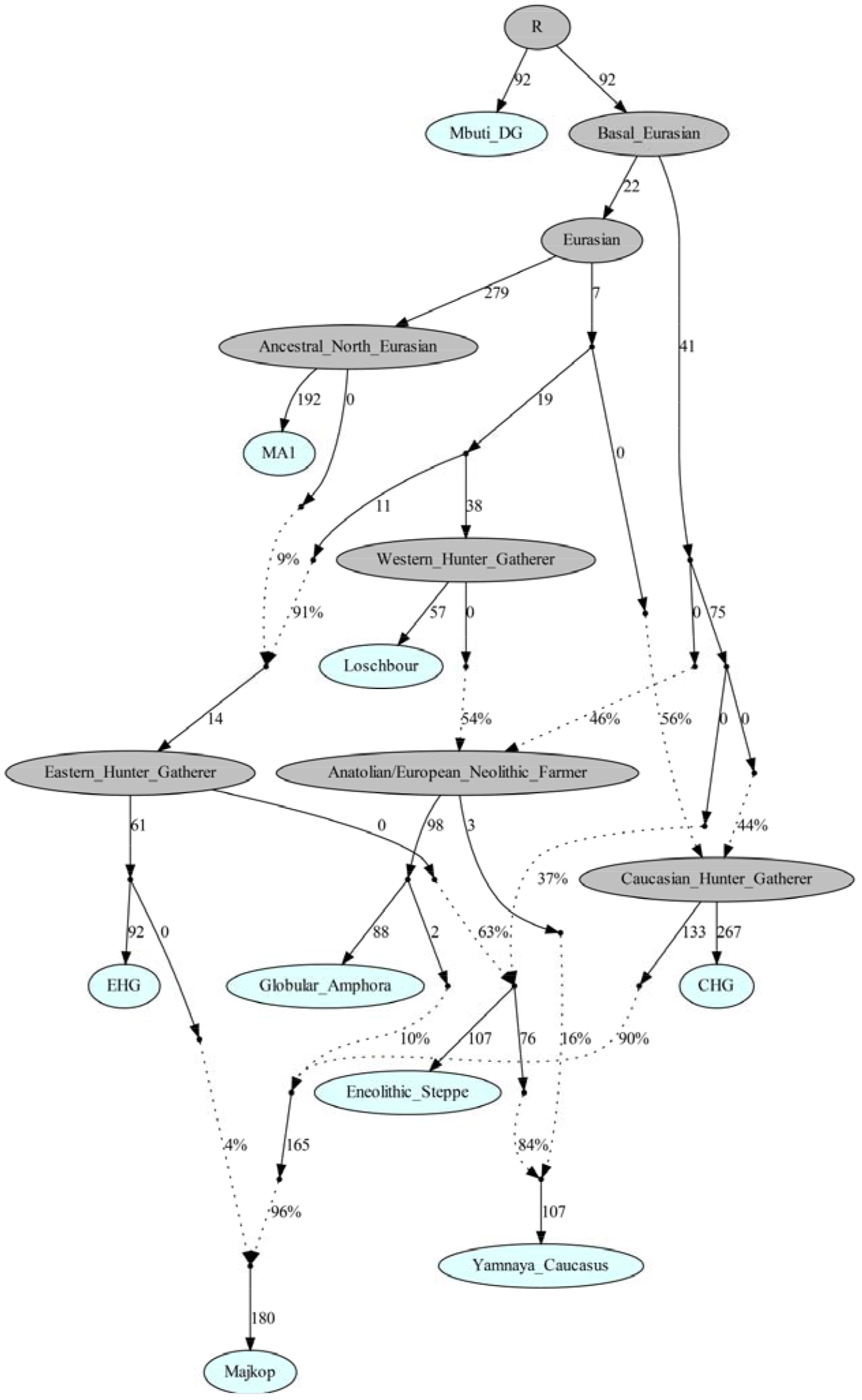
Admixture Graph modelling of the population history of the Caucasus region. We started with a skeleton tree without admixture including Mbuti, Loschbour and MA1. We grafted onto this EHG, CHG, Globular_Amphora, Eneolithic_steppe, Maykop, and Yamnaya_Caucasus, adding them consecutively to all possible edges in the tree and retaining only graph solutions that provided no differences of |Z|>3 between fitted and estimated statistics. The worst match is |Z|=2.824 for this graph. We note that the maximum discrepancy is *f_4_*(MA1, Maykop; EHG, Eneolithic_steppe)=-3.369 if we do not add the 4% EHG ancestry to Maykop. Drifts along edges are multiplied by 1000 and dashed lines represent admixture.

## Discussion

Our data from the Greater Caucasus region cover over 3000 years of prehistory as a transect through time, ranging from the Eneolithic (starting 6500 yBP, 4500 calBCE) to the Late Bronze Age (ending 3200 yBP, 1200 calBCE). We observe a genetic separation between the groups in the piedmont steppe, i.e. the northern foothills of the Greater Caucasus, and those groups of the bordering herb, grass and desert steppe regions in the north (i.e. the ‘real’ steppe). We have summarised these broadly as *Caucasus* and *Steppe* groups in correspondence with the eco-geographic vegetation zones that characterise the socio-economic basis of the associated archaeological cultures.

When compared to present-day human populations from the Caucasus, which show a clear separation into North and South Caucasus groups along the Great Caucasus mountain range (Fig. 2D), our new data highlights that the situation during the Bronze Age was quite different. The fact that individuals buried in kurgans in the North Caucasian piedmont and foothill zone are more closely related to ancient individuals from regions further south in today’s Armenia, Georgia and Iran allows us to draw two major conclusions.

First, sometime after the Bronze Age present-day North Caucasian populations must have received additional gene-flow from populations north of the mountain range that separates them from southern Caucasians, who largely retained the Bronze Age ancestry profile. The archaeological and historic records suggest numerous incursions during the subsequent Iron Age and Medieval times^48^, but ancient DNA from these time periods is needed to test this directly.

Second, our results reveal that the Greater Caucasus Mountains were not an insurmountable barrier to human movement in prehistory. Instead the foothills to the north at the interface of the steppe and mountain ecozones could be seen as a transfer zone of cultural innovations from the south and the adjacent Eurasian steppes to the north, as attested by the archaeological record. The latter is best exemplified by the two Steppe Maykop outlier individuals dating to 5100-5000 yBP/3100-3000 calBCE, which carry additional Anatolian farmer-related ancestry likely derived from a proximate source related to the *Caucasus* cluster. We could show that individuals from the contemporaneous Maykop period in the piedmont region are likely candidates for the source of this ancestry and might explain the regular presence of ‘Maykop artefacts’ in burials that share Steppe Eneolithic traditions and are genetically assigned to the *Steppe* group. Hence the diverse ‘Steppe Maykop’ group indeed represents the mutual entanglement of *Steppe* and *Caucasus* groups and their cultural affiliations in this interaction sphere.

Concerning the influences from the south, our oldest dates from the immediate Maykop predecessors Darkveti-Meshoko (Eneolithic Caucasus) indicate that the *Caucasus* genetic profile was present north of the range ~6500 BP, 4500 calBCE.This is in accordance with the Neolithization of the Caucasus, which had started in the flood plains of the great rivers in the South Caucasus in the 6^th^ millennium BCE from where it spread to the West and Northwest Caucasus during the 5^th^ millennium BCE^9,49^. It remains unclear whether the local CHG ancestry profile (represented by Late Upper Palaeolithic/Mesolithic individuals from Kotias Klde and Satsurblia in today’s Georgia) was also present in the North Caucasus region before the Neolithic.

However, if we take the Caucasus hunter-gatherer individuals from Georgia as a local baseline and the oldest Eneolithic Caucasus individuals from our transect as a proxy for the local Late Neolithic ancestry, we notice a substantial increase in Anatolian farmer-related ancestry. This in all likelihood is linked to the process of Neolithization, which also brought this type of ancestry to Europe. As a consequence, it is possible that Neolithic groups could have reached the northern flanks of the Caucasus earlier^50^ (Supplementary Information 1) and in contact with local hunter-gatherers facilitated the exploration of the steppe environment for pastoralist economies. Hence, additional sampling from older individuals is needed to fill this temporal and spatial gap.

Our results show that at the time of the eponymous grave mound of Maykop, the North Caucasus piedmont region was genetically connected to the south. Even without direct ancient DNA data from northern Mesopotamia, the new genetic evidence suggests an increased assimilation of Chalcolithic individuals from Iran, Anatolia and Armenia and those of the Eneolithic Caucasus during 6000-4000 calBCE^23^, and thus likely also intensified cultural connections. Within this sphere of interaction, it is possible that cultural influences and continuous subtle gene flow from the south formed the basis of Maykop (Fig. 4; Supplementary Table 10). In fact, the Maykop phenomenon was long understood as the terminus of the expansion of South Mesopotamian civilisations in the 4^th^ millennium BCE^11, 12, 51^. It has been further suggested that along with the cultural and demographic influence the key technological innovations that had revolutionised the late 4^th^ millennium BCE in western Asia had ultimately also spread to Europe^52^. An earlier connection in the late 5^th^ millennium BCE, however, allows speculations about an alternative archaeological scenario: was the cultural exchange mutual and did e.g. metal rich areas such as the Caucasus contribute substantially to the development and transfer of these innovations^53, 54^?

We also observe a degree of genetic continuity within each cluster. While this continuity in each cluster spans the 3000 years covered in this study, we also detect occasional gene-flow between the two clusters as well as from outside sources. Moreover, our data shows that the northern flanks were consistently linked to the Near East and had received multiple streams of gene flow from the south, as seen e.g. during the Maykop, Kura-Araxes and late phase of the North Caucasus culture. Interestingly, this renewed appearance of the southern genetic make-up in the foothills corresponds to a period of climatic deterioration (known as 4.2 ky event) in the steppe zone, that put a halt to the exploitation of the steppe zone for several hundred years^55^. Further insight arises from individuals that were buried in the same kurgan but in different time periods, as highlighted in the two kurgans Marinskaya 5 and Sharakhalsun 6. Here, we recognize that the distinction between *Steppe* and *Caucasus* with reference to vegetation zones(Fig. 1) is not strict but rather reflects a shifting border of genetic ancestry through time, possibly due to climatic shifts and/or cultural factors linked to subsistence strategies or social exchange. It seems plausible that the occurrence of *Steppe* ancestry in the piedmont region of the northern foothills coincides with the range expansion of the Yamnaya pastoralists. However, more time-stamped data from this region will be needed to provide further details on the dynamics of this contact zone.

An interesting observation is that steppe zone individuals directly north of the Caucasus (Eneolithic Samara and Eneolithic steppe) had initially not received any gene flow from Anatolian farmers. Instead, the ancestry profile in Eneolithic steppe individuals shows an even mixture of EHG and CHG ancestry, which argues for an effective cultural and genetic border between the contemporaneous Eneolithic populations in the North Caucasus, notably *Steppe* and *Caucasus.* Due to the temporal limitations of our dataset, we currently cannot determine whether this ancestry is stemming from an existing natural genetic gradient running from EHG far to the north to CHG/Iran in the south or whether this is the result of farmers with Iranian farmer/ CHG-related ancestry reaching the steppe zone independent of and prior to a stream of Anatolian farmer-like ancestry, where they mixed with local hunter-gatherers that carried only EHG ancestry.

Another important observation is that all later individuals in the steppe region, starting with Yamnaya, deviate from the EHG-CHG admixture cline towards European populations in the West. This documents that these individuals had received Anatolian farmer-related ancestry, as documented by quantitative tests and recently also shown for two Yamnaya individuals from Ukraine (Ozera) and one from Bulgaria^24^. For the North Caucasus region, this genetic contribution could have occurred through immediate contact with groups in the *Caucasus* or further south. An alternative source, explaining the increase in WHG-related ancestry, would be contact with contemporaneous Chalcolithic/EBA farming groups at the western periphery of the Yamnaya culture distribution area, such as Globular Amphora and Tripolye (Cucuteni–Trypillia) individuals from Ukraine, which also have been shown to carry Anatolian Neolithic farmer-derived ancestry^24^.

Archaeological arguments would be consonant with both scenarios. Contact between early Yamnaya and late Maykop groups at the end of the 4^th^ millennium BCE is suggested by impulses seen in early Yamnaya complexes. A western sphere of interaction is evident from striking resemblances of imagery inside burial chambers of Central Europe and the Caucasus^56^ (Supplementary Fig. 9), and particular similarities also exist in geometric decoration patterns in stone cist graves in the Northern Pontic steppe^57^, on stone *stelae* in the Caucasus^58^, and on pottery of the Eastern Globular Amphora Culture, which links the eastern fringe of the Carpathians and the Baltic Sea^56^. This implies an overlap of symbols with a communication and interaction network that formed during the late 4^th^ millennium BCE and operated across the Black Sea area involving the Caucasus^59, 60^, and later also involved early Globular Amphora groups in the Carpathians and east/central Europe^61^. The role of early Yamnaya groups within this network is still unclear^57^. However, this interaction zone pre-dates any direct influence of Yamnaya groups in Europe or the succeeding formation of the Corded Ware^62, 63^ and its persistence opens the possibility of subtle bidirectional gene-flow, several centuries before the massive range expansions of pastoralist groups that reached Central Europe in the mid-3^rd^ millennium BCE^19, 35^.

We were surprised to discover that Steppe Maykop individuals from the eastern desert steppes harboured a distinctive ancestry component that relates them to Upper Palaeolithic Siberian individuals (AG3, MA1) and Native Americans. This is exemplified by the more commonly East Asian features such as the derived EDAR allele, which has also been observed in EHG from Karelia and Scandinavian hunter-gatherers (SHG). The additional affinity to East Asians suggests that this ancestry does not derive directly from Ancestral North Eurasians but from a yet-to-be-identified ancestral population in north-central Eurasia with a wide distribution between the Caucasus, the Ural Mountains and the Pacific coast^21^, of which we have discovered the so far southwestern-most and also youngest (e.g. the Lola culture individual) genetic representative.

The insight that the Caucasus mountains served not only as a corridor for the spread of CHG/Neolithic Iranian ancestry but also for later gene-flow from the south also has a bearing on the postulated homelands of Proto-Indo-European (PIE) languages and documented gene-flows that could have carried a consecutive spread of both across West Eurasia^17, 64^. Perceiving the Caucasus as an occasional bridge rather than a strict border during the Eneolithic and Bronze Age opens up the possibility of a homeland of PIE south of the Caucasus, which itself provides a parsimonious explanation for an early branching off of Anatolian languages. Geographically this would also work for Armenian and Greek, for which genetic data also supports an eastern influence from Anatolia or the southern Caucasus. A potential offshoot of the Indo-Iranian branch to the east is possible, but the latest ancient DNA results from South Asia also lend weight to an LMBA spread via the steppe belt^21^.The spread of some or all of the proto-Indo-European branches would have been possible via the North Caucasus and Pontic region and from there, along with pastoralist expansions, to the heart of Europe. This scenario finds support from the well attested and now widely documented ‘steppe ancestry’ in European populations, the postulate of increasingly patrilinear societies in the wake of these expansions (exemplified by R1a/R1b), as attested in the latest study on the Bell Beaker phenomenon^35^.

## Materials and Methods

### Sample collection

Samples from archaeological human remains were collected and exported under a collaborative research agreement between the Max-Planck Institute for the Science of Human History, the German Archaeological Institute and the Lomonosov Moscow State University and Anuchin Research Institute and Museum of Anthropology (permission no. № 114-18/204-03).

### Ancient DNA analysis

We extracted DNA and prepared next-generation sequencing libraries from 107 samples in two dedicated ancient DNA laboratories at Jena and Boston. Samples passing initial QC were further processed at the Max Planck Institute for the Science of Human History, Jena, Germany following the established protocols for DNA extraction and library preparation^65, 66^. Fourteen of these samples were processed at Harvard Medical School, Boston, USA following a published protocol by replacing the extender-MinElute-column assembly with the columns from the Roche High Pure Viral Nucleic Acid Large Volume Kit to extract DNA from about 75mg of sample powder from each sample. All libraries were subjected to partial (“half”) Uracil-DNA-glycosylase (UDG) treatment before blunt end repair. We performed in-solution enrichment (1240K capture)^27^ for a targeted set of 1,237,207 SNPs that comprises two previously reported sets of 394,577 SNPs (390k capture) and 842,630 SNPs, and then sequenced on an in-house Illumina HiSeq 4000 or NextSeq 500 platform for 76bp either single or paired-end.

The sequence data was demultiplexed, adaptor clipped with leehom^67^ and then further processed using *EAGER*^68^, which included mapping with *BWA (v0.6.l)*^69^ against human genome reference GRCh37/hg19, and removing duplicate reads with the same orientation and start and end positions. To avoid an excess of remaining C-to-T and G-to-A transitions at the ends of the reads, three bases of the ends of each read were clipped for each sample using trimBam (https://genome.sph.umich.edu/wiki/BamUtil:_trimBam). We generated “pseudo-haploid” calls by selecting a single read randomly for each individual at each of the targeted SNP positions using the in-house genotype caller *pileupCaller* (https://github.com/stschiff/sequenceTools/tree/master/src-pileupCaller).

### Quality control

We report, but have not analyzed, data from individuals that had less than 30,000 SNPs hit on the 1240K set. We removed individuals with evidence of contamination based on heterozygosity in the mtDNA genome data, a high rate of heterozygosity on the X chromosome despite being male estimated with *ANGSD*^70^, or an atypical ratio of the reads mapped to X versus Y chromosomes.

### Merging new and published ancient and modern population data

We merged our newly generated ancient samples with ancient populations from the publicly available datasets^13, 19, 20, 24, 27, 28, 33, 35, 37^ (Supplementary Data 2), as well as genotyping data from worldwide modern populations using Human Origins arrays published in the same publications. We also included newly genotyped populations from the Caucasus and Asia, described in detail in Jeong et al.^71^.

### Principal Component Analysis

We carried out principal component analysis on Human Origins Dataset using the *smartpca* program of *EIGENSOFT*^44^, using default parameters and the lsqproject: YES, numoutlieriter: 0, and shrinkmode:YES options to project ancient individuals onto the first two components.

### ADMIXTURE analysis

We carried out *ADMIXTURE (v1.23)^45^* analysis after pruning for linkage disequilibrium in *PLINK^72^* with parameters --indep-pairwise 200 25 0.4, which retained 301,801 SNPs for the Human Origins Dataset. We ran *ADMIXTURE* with default 5-fold cross-validation (--cv=5), varying the number of ancestral populations between K=2 and K=22 in 100 bootstraps with different random seeds.

### *f*-statistics

We computed *D*-statistics and *f*_4_-statistics using *qpDstat* program of *ADMIXTOOLS^44^* with default parameters. We computed the admixture *f_3_*-statistics using the *qp3Pop* program of *ADMIXTOOLS* with the flag inbreed: YES. *ADMIXTOOLS* computes standard errors using the default block jackknife.

### Testing for streams of ancestry and inference of mixture proportions

We used *qpWave* and *qpAdm*^19^ as implemented in *ADMIXTOOLS* to test whether a set of test populations is consistent with being related via *N* streams of ancestry from a set of outgroup populations and estimate mixture proportions for a *Test* population as a combination of *N*’reference’ populations by exploiting (but not explicitly modeling) shared genetic drift with a set of outgroup populations. Mbuti.DG, Ust_Ishim.DG, Kostenki14, MA1, Han.DG, Papuan.DG, Onge.DG, Villabruna, Vestonice16, ElMiron, Ethiopia_4500BP.SG, Karitiana.DG, Natufian, Iran_Ganj_Dareh_Neolithic. The “DG” samples are extracted from high coverage genomes sequenced as part of the Simons Genome Diversity Project^33^. For some analyses, we used an extended set of outgroup populations, including some of the following additional ancient populations to constrain standard errors: WHG, EHG, and Levant Neolithic.

### Dating of gene-flow events

We estimated the time depth of selected admixture events using the linkage disequilibrium (LD)-based admixture inference implemented in *ALDER*^46^.

### Admixture graph modelling

Admixture graph modelling was carried out with the *qpGraph* software as implemented in *ADMIXTOOLS*^44^ using Mbuti.DG as an outgroup.

### Sex determination and Y chromosomal and mtDNA haplogroup assignment

We determined the sex of the newly reported samples in this study by counting the number of reads overlapping with the targets of 1240k capture reagent^37^. We extracted the reads of high base and mapping quality (samtools depth -q30 -Q37) using *samtools v 1.3.1*^73^. We calculated the ratios of the numbers of reads mapped on X chromosome or Y chromosome compared with that mapped on autosomes (X-rate and Y-rate, respectively). Samples with an X-rate<0.42 and a Y-rate>0.26 were assigned as males and those with an X-rate>0.68 and a Y-rate<0.02 were assigned as females.

We used *EAGER* and *samtools v1.3.1* to extract reads from the 1240k SNP and mitocapture data mapped to the rCRS. We used *GeneiousR8.1.9*^74^ to locally realign, visually inspect the pileups for contamination, and to call consensus sequences, which were used for haplotyping in *HaploGrep 2*^75^. In addition, we used the software contamMix 1.0.10, which employs a Bayesian approach to estimate contamination in the mitochondrial genome^76^.

We called Y chromosomal haplogroups for males using the captured SNPs on Y chromosome by restricting to sequences with mapping quality ≥30 and bases with base quality ≥30. We determined Y chromosomal haplogroups by identifying the most derived allele upstream and the most ancestral allele downstream in the phylogenetic tree in the ISOGG version 11.89 (accessed March 31, 2016) (http://www.isogg.org/tree).

### Kinship analysis

We used outgroup-*f_3_* statistics and the methods *lcMLkin^77^* and *READ^78^* to determine genetic kinship between individuals.

### Phenotypic SNP calls

We determined the allele information of 5 SNPs (rs4988235, rs16891982, rs1426654, rs3827760, rs12913832) thought to be affected by selection in our ancient samples using the captured SNPs by restricting to sequences with mapping quality ≥30 and bases with base quality ≥30 (Supplementary Information 7).

### Abbreviations

We use the following abbreviated labels throughout the manuscript: E, Early; M, Middle; L, Late; N, Neolithic; BA, Bronze Age; WHG, EHG, CHG, Western,
Eastern, Caucasus hunter-gatherers, respectively; Mal’ta 1, MA1; Afontova Gora 3, AG3.

### Data availability

Data is deposited in the European Nucleotide Archive under the accession numbers XXX-XXX (will be made available during revision).

## Acknowledgments

We thank Stephen Clayton and all members of the MPI-SHH Archaeogenetics Department for support, Michelle O’Reilly and Hans Sell for graphics support, and Iosif Lazaridis and Nick Patterson for critical discussions. We thank Susanne Lindauer, Ronny Friedrich, Robin van Gyseghem and Ute Blach for radiocarbon dating support. This work was funded by the Max Planck Society and the German Archaeological Institute (DAI). C.C.W. was funded by Nanqiang Outstanding Young Talents Program of Xiamen University (X2123302) and the Fundamental Research Funds for the Central Universities.

## Author contributions

SH, JK, CCW, SR and WH conceived the idea for the study design. AW, GB, OC, MF, EH,DK, SM, NR, KS and WH performed and supervised wet and dry lab work. SH, AK, ARK, VEM,VGP, VRE, BCA, RGM, PLK, KWA, SLP, CG, HM, BV, LY, ADR, DM, NYB, JG, KF, CK, YBB, APB,VT, RP, SH and ABB assembled skeletal material, contextual information and provided site descriptions. CCW, SR and WH analysed data. CJ, IM, SS, EB, OB provided additional data and methods. WH, CCW, SR, SH, VT, RP, TH, DR and JK wrote the manuscript with input from all authors.

